# GCRTcall: a Transformer based basecaller for nanopore RNA sequencing enhanced by gated convolution and relative position embedding via joint loss training

**DOI:** 10.1101/2024.06.03.597255

**Authors:** Qingwen Li, Chen Sun, Daqian Wang, Jizhong Lou

## Abstract

Nanopore sequencing, renowned for its ability to sequence DNA and RNA directly with read lengths extending to several hundred kilobases or even megabases, holds significant promise in fields like transcriptomics and other omics studies. Despite its potential, the technology’s limited accuracy in base identification has restricted its widespread application. Although many algorithms have been developed to improve DNA decoding, advancements in RNA sequencing remain limited. Addressing this challenge, we introduce GCRTcall, a novel approach integrating Transformer architecture with gated convolutional networks and relative positional encoding for RNA sequencing signal decoding. Our evaluation demonstrates that GCRTcall achieves state-of-the-art performance in RNA basecalling.

## Introduction

Nanopore sequencing technology directly sequences single strands of DNA or RNA by detecting changes in electrical current as the molecules pass through nanopores, eliminating the need for PCR amplification. This technique enables rapid single-molecule sequencing with significantly increased read lengths, reaching hundreds of kilobases or even magabases. It holds immense potential in various omics sequencing studies such as genomics, transcriptomics, epigenomics, and proteomics ^[1-11]^.

Despite its advantages, the accuracy of basecalling has emerged as a significant bottleneck, limiting further broader application of nanopore sequencing. Sequencing signals are influenced not only by individual nucleotides but also by neighboring bases, resulting in non-uniform translocation of the sequences and low signal-to-noise ratios measured in picoamperes (pA). These challenges make accurate basecalling in nanopore sequencing particularly difficult ^[3, 12, 13]^.

In recent years, several algorithms have been developed to improve the accuracy of nanopore sequencing signal decoding. Methods like Metrichor and Nanocall^[14]^, which utilize Hidden Markov Models (HMM), segment events in the current signal and calculate transition probabilities for basecalling. Other approaches, such as Chiron^[15]^, Deepnano^[16]^, and Guppy, leverage Recurrent Neural Network (RNN) architectures, while Causalcall^[17]^ and RODAN^[18]^ employ Convolutional Neural Network (CNN) architectures to achieve end-to-end basecalling. Additionally, SACall incorporates self-attention mechanisms into nanopore signal decoding^[19]^. However, with the exception of RODAN^[18]^, the focus of these methods is primarily on DNA basecalling, with limited research in RNA decoding.

Unlike several hundreds base pairs per second (bps) translocation speed for DNA, RNA translocates at only about or below 100 bps. Additionally, there are substantial differences in the physical and chemical properties between DNA and RNA, resulting in distinct signal patterns. Consequently, models designed for DNA basecalling are usually ineffective for RNA signal decoding. To address this gap, we propose GCRTcall, a **T**ransformer based base**call**er for nanopore RNA sequencing, enhanced by **G**ated **C**onvolution and **R**elative position embedding through joint loss training. This method achieves state-of-the-art decoding accuracy on multi-species transcriptome sequencing data.

## Materials and Methods

### Benchmark Dataset

The benchmark dataset used in this study was proposed by Neumann et al. ^[18]^, which is also utilized in the development of RODAN^[18]^. The training set comprises five species: Arabidopsis thaliana^[18]^, Epinano synthetic constructs^[20]^, Homo sapiens^[21]^, Caenorhabditis elegans^[22]^, and Escherichia coli^[23]^. Initially, all reads were basecalled using Guppy version 6.2.1^[24]^. The decoded sequences were then mapped to the reference genome with minimap2 ^[25]^ to obtain corrected sequences. Subsequently, Taiyaki ^[26]^ was utilized to generate an HDF5 file containing the raw signal of each read, its corresponding corrected sequence, and their mapping relationship. The training dataset contained 116,072 reads: with 24,370 from Arabidopsis, 29,728 from Epinano synthetic constructs, 30,048 from H. sapiens, 24,192 from C. elegans, and 7,734 from E. coli. The test set included also five species: Homo sapiens^[21]^, Arabidopsis thaliana^[27]^, Mus musculus^[28]^, S. cerevisiae^[29]^, and Populus trichocarpa^[30]^, each consisting of 100,000 reads.

### Model Architecture

Our model architecture was inspired by Google’s Conformer^[31]^, a convolution-augmented Transformer known for effectively modeling both global and local dependencies, outperforming traditional Transformer^[32, 33]^ and CNN^[34-37]^ models in speech recognition tasks. GCRTcall compreises three CNN layers for downsampling and feature extraction, with output channels of 4, 6, and 512, and convolutional kernels of size 5, 5, and 19, with strides of 1, 1, and 10, respectively. This is followed by 8 Conformer blocks and a connectionist temporal classification (CTC) decoder ^[31]^, amounting to a total of 50 million parameters. Our previous study indicated that training with a joint loss, combining CTC loss and Kullback-Leibler Divergence (KLDiv) loss, results in superior basecalling accuracy compared to using only CTC loss under the same inference structure^[38]^. Therefore, during the training phase, GCRTcall incorporates additional forward and backward Transformer decoders at the top, utilizing the joint loss for improved convergence. The model architecture of GCRTcall is illustrated in Figure 1.

**Figure 1.**
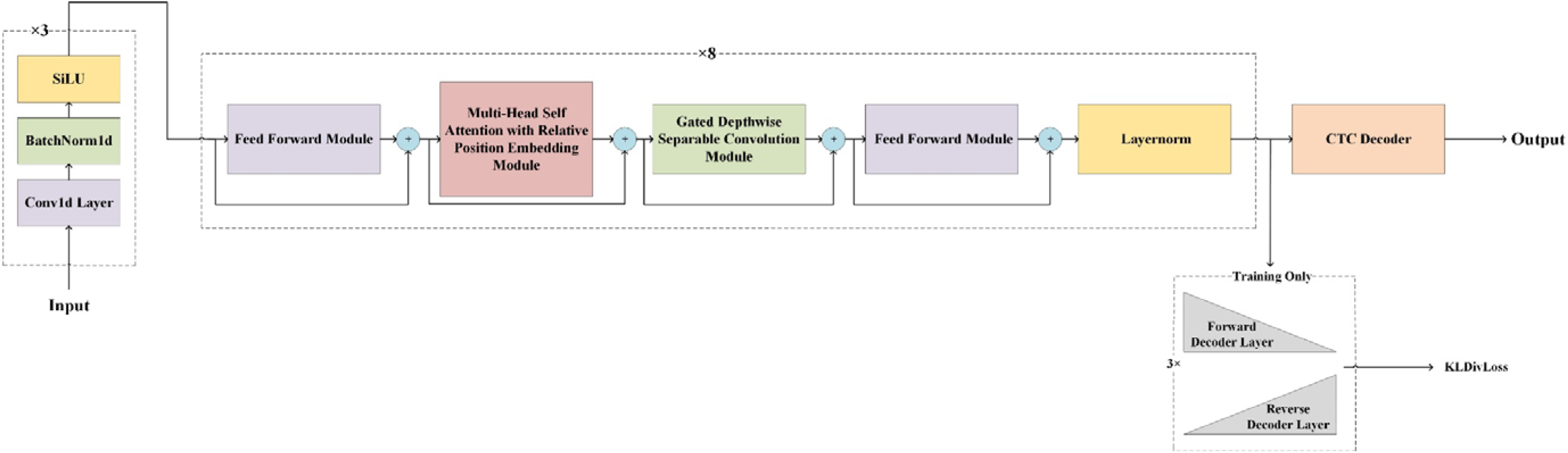
Schematics representation of the architecture of GCRTcall. GCRTcall compreises three CNN layers for downsampling and feature extraction, and followed by 8 Conformer blocks and a CTC decoder. During training, a pair of forward and reverse decoder was added on top of the base architecture for joint loss training.

Compared to traditional Transformers, the Conformer modules in GCRTcall feature two key improvements: First, they combine relative positional embedding with a multi-head self-attention mechanism to enhance the model’s robustness to inputs of varying lengths. Second, they integrate depthwise separable convolutions based on gate mechanisms to process the outputs of attention layers, thereby strengthening the model’s ability to capture local dependencies within sequences.

### Relative Position Multi-Head Self-Attention Mechanism

Transformer-XL^[39]^ integrates relative positional embedding with a self-attention mechanism, enhancing the model’s representational capacity for sequences of varying lengths. The relative position multi-head self-attention mechanism processes input sequences along with its sinusoidal position encoding. It performs three linear projections on the input to generate *Q, K*, and *V*, and also applying linear projection on positional embedding to obtain *K*p. Two biases, *bk* and *bp*, are initialized. The computation principle of the relative position self-attention mechanism is as follows:

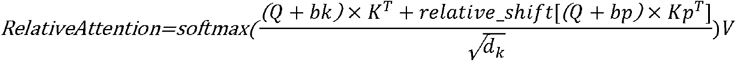

The multi-head attention then combines and projects the aforementioned attention computation results as follows:

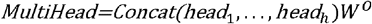

### Gated Depthwise Separable Convolution

EfficientNet^[40]^ utilizes depthwise separable convolution to reduce the number of parameters and enhance computational efficiency while maintaining state-of-the-art accuracy. Similarly, Dauphin et al.^[41]^ proposed gated convolutional networks, which utilize CNNs to extract hidden states from sequences and employ gated linear units (GLU) to augment non-linear expression and mitigate the vanishing gradient problem. This approach enables the model to compute in parallel, outperforming LSTMs on multiple NLP datasets. GLU is computed as follows:

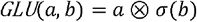

where *a* is the first half of the input matrices and *b* is the second half.

Inspired by these approaches, the structure of the gated depthwise separable convolution block in GCRTcall is illustrated in Figure 2.

**Figure 2.**
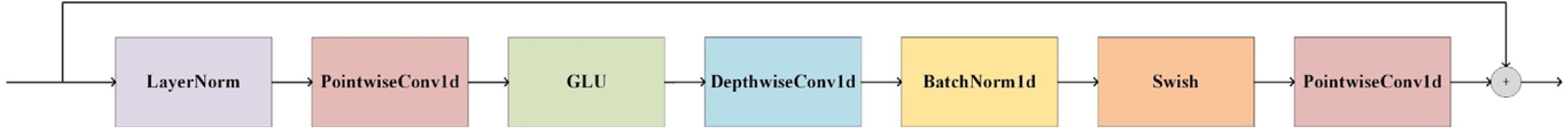
Architecture of the gated depthwise separable convolution block of GCRTcall. The convolution block consists of a 1-D pointwise convolution followed by a GLU, a 1-D depthwise convolution, 1-D batch normalization, and a swish activation function.

### Joint loss training

An additional forward and reverse transformer decoder were added on the top of the inference structure of the model during training. The forward decoder adopts a lower triangular matrix as a causal mask, while the reverse decoder is equipped with an anti-lower triangular causal mask.

The model is trained by optimizing a joint loss that includes CTC loss and KLDiv loss to ensure convergence.

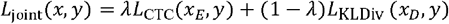

Where *x*_E_ is the output probability matrix of the encoder, and *x*_D_ is the output of the decoders, *y* is the label, λ is a hyperparameter between 0 and 1. In this paper, λ was set to 0.5.

### Model Training

As previously demonstrated, using a joint loss that combines CTC loss and KLdiv loss can help accelerate model convergenc^[38]^. Therefore, during training, we added two layers of forward and backward Transformer decoders at the top of GCRTcall, which are not utilized during actual inference.

GCRTcall was trained on an Ubuntu server equipped with 2×2.10GHz 36-core CPUS, providing 144 logical CPUs and 512GB of RAM. The training utilized 2 NVIDIA RTX 6000 Ada Generation 48G GPUs for 12 epochs, totaling 12.95 hours. The batch size of 140 was employed, managed by the Ranger optimizer at a learning rate of 0.002. The training was conducted using the ReduceLROnPlateau learning rate scheduler based on validation set loss monitoring.

### Performance Evaluation

Identity, mismatch rate, insertion rate, and deletion rate were adopted as metrics to evaluate the decoding accuracy of the model. These overall median metrics are commonly used in multiple basecaller researches for performance evaluation and comparation:

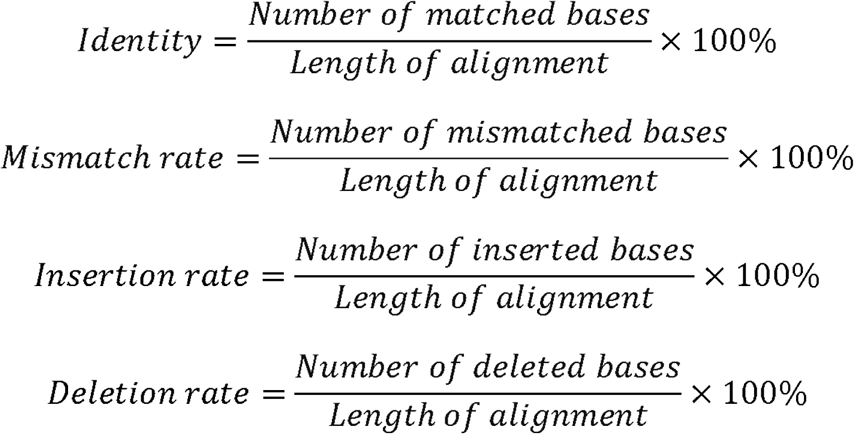

## Results and Discussion

### Comparison of Decoding Performance with Different Basecallers

We compared the basecalling accuracy of GCRTcall, Guppy 6.2.1, and RODAN on a test set consisting of five species. All basecalling results were aligned to the reference genomes using minimap2, retaining only the optimal alignment results. As shown in Table 1, GCRTcall achieved state-of-the-art accuracy levels across all five species. Notably, according to Neumann et al.^[18]^, while RODAN slightly outperforms Guppy in basecalling accuracy for mouse and yeast, GCRTcall significantly outperforms both in decoding accuracy for these two species. Additionally, all three basecallers exhibited the poorest performance on mouse, consistent with previous findings that suggest substantial differences in sequencing signal patterns between mice and other species.

**Table 1.** Performance comparation between GCRTcall, RODAN, and Guppy.

| Species | Basecaller | Identity (%) | Insertion rate (%) | Deletion rate (%) | Mismatch rate (%) |
| --- | --- | --- | --- | --- | --- |
| Yeast | Guppy | 91.36 | 1.57 | 2.48 | 4.56 |
|  | RODAN | 90.86 | 1.67 | 2.52 | 5.17 |
|  | GCRTcall | **92.59** | **1.54** | **2.02** | **3.83** |
| Human | Guppy | 90.59 | 1.42 | 3.02 | 4.89 |
|  | RODAN | 92.75 | 1.27 | 2.09 | 3.69 |
|  | GCRTcall | **94.22** | **1.10** | **1.68** | **2.85** |
| Mouse | Guppy | 87.67 | 1.73 | 3.70 | 6.84 |
|  | RODAN | 86.82 | **1.41** | 3.97 | 7.36 |
|  | GCRTcall | **89.96** | 1.44 | **3.07** | **5.44** |
| Poplar | Guppy | 90.15 | 1.66 | 2.81 | 5.31 |
|  | RODAN | 90.55 | **1.49** | 2.53 | 5.36 |
|  | GCRTcall | **92.25** | 1.51 | **2.07** | **4.08** |
| Arabidopsis | Guppy | 91.58 | 1.33 | 2.69 | 4.36 |
|  | RODAN | 93.93 | **1.10** | 1.87 | 2.63 |
|  | GCRTcall | **94.03** | 1.14 | **1.77** | **2.94** |

The inference was conducted on an Ubuntu server equipped with an Intel i9-13900K CPU, 125G RAM, and one NVIDIA RTX 3090 24G GPU. The inference speed of different basecaller model was also evaluated and compared. A random read with 37,796 sampling points was selected as input to test the inference time of each model. Guppy achieved the fastest decoding speed at 1.02E+07 samples per second, owing to its smaller parameter count of 2.2M. RODAN followed at 4.68E+06 samples per second, while GCRTcall, with its 50M parameters, completed decoding with speed at 1.68E+06 samples per second. Several acceleration optimization algorithms for Transformer-based models, such as hardware-aware techniques, sparse attention, and model quantization, have been proposed to enhance inference speed. These algorithms will be tested in the future development of GCRTcall.

### Ablation Study

To further explore the impact of model structures on the basecalling accuracy of GCRTcall, we conducted two sets of ablation experiments: first, removing relative shift operation for position scores (GCRTcall w/o RS); and second, replacing Conformer modules with Transformer modules (Transcall).

In Transformer-XL, absolute position representation is initially performed to reduce the computational complexity of relative positional encoding. A relative shift of position scores is then applied to obtain relative position embeddings for sequences. To investigate the impact of relative position embeddings on model performance, GCRTcall was trained without the relative shift operation for 12 epochs using the same training set. The test results (Table 2) show a decrease in decoding performance compared to GCRTcall, indicating that relative position embeddings enhance the robustness of attention mechanisms to sequence position representation. To investigate the impact of gated convolution neural networks on model performance, Transformer modules were used to replace Conformer modules, and the model was also trained for 12 epochs. The test results (Table 2) indicate that the model’s decoding performance deteriorated compared to GCRTcall and GCRTcall w/o RS. This suggests that gated convolutional networks, which enhance the representation of local dependencies, play an important role in accurate basecalling.

**Table 2.** Identity comparation between GCRTcall, GCRTcall w/o RS, and Transcall.

| Species | Basecaller | Identity (%) | Insertion rate (%) | Deletion rate (%) | Mismatch rate (%) |
| --- | --- | --- | --- | --- | --- |
| Yeast | Transcall | 90.07 | 1.83 | 2.98 | 5.07 |
|  | GCRTcall w/o RS | 91.54 | 1.65 | 2.44 | 4.30 |
|  | GCRTcall | **92.59** | **1.54** | **2.02** | **3.83** |
| Human | Transcall | 91.24 | 1.39 | 2.97 | 4.28 |
|  | GCRTcall w/o RS | 93.03 | 1.33 | 2.02 | 3.38 |
|  | GCRTcall | **94.22** | **1.10** | **1.68** | **2.85** |
| Mouse | Transcall | 87.02 | 1.49 | 4.46 | 6.93 |
|  | GCRTcall w/o RS | 88.61 | 1.61 | 3.57 | 6.11 |
|  | GCRTcall | **89.96** | **1.44** | **3.07** | **5.44** |
| Poplar | Transcall | 89.71 | 1.76 | 3.14 | 5.30 |
|  | GCRTcall w/o RS | 91.06 | 1.69 | 2.47 | 4.62 |
|  | GCRTcall | **92.25** | **1.51** | **2.07** | **4.08** |
| Arabidopsis | Transcall | 91.24 | 1.40 | 2.97 | 4.28 |
|  | GCRTcall w/o RS | 92.87 | 1.31 | 2.19 | 3.45 |
|  | GCRTcall | **94.03** | **1.14** | **1.77** | **2.94** |

The training curves of GCRTcall, GCRTcall w/o RS, and Transcall are illustrated in Figure 3. It can be observed that the form of position encoding has little impact on convergence during training, mainly enhancing the model’s generalization ability for decoding sequences of varying lengths. However, Transcall, without convolutional enhancement, converges slower and to a higher loss compared to both GCRTcall and GCRTcall w/o RS.

**Figure 3.**
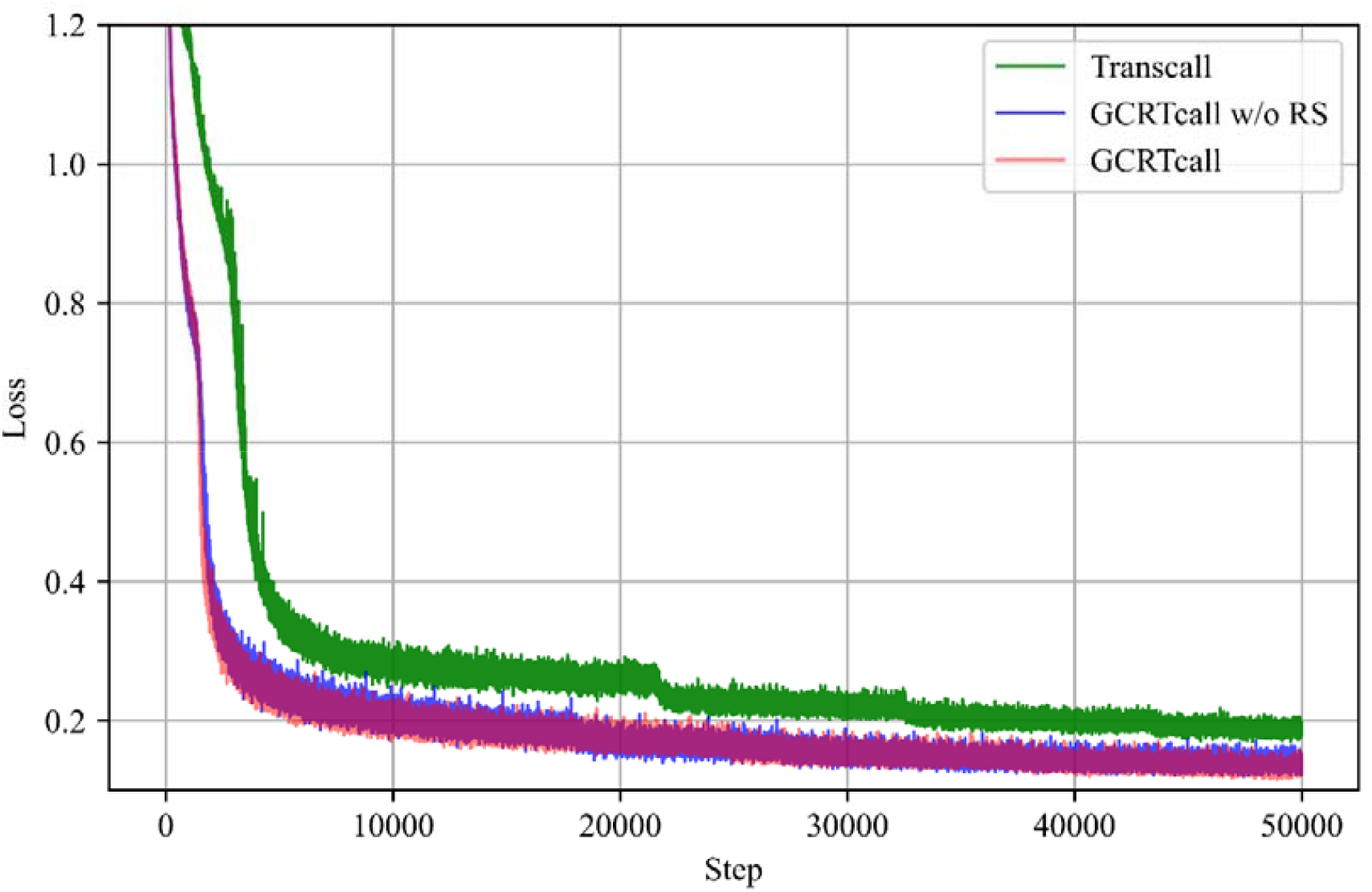
Training curves of GCRTcall, GCRTcall w/o RS, and Transcall. GCRTcall and GCRTcall w/o RS exhibit similar training curve. While Transcall, without convolutional enhancement, converges slower and to a higher loss compared to both GCRTcall and GCRTcall w/o RS.

## Conclusion

This study introduces GCRTcall, a Transformer-based basecaller designed for nanopore RNA sequencing signal decoding. GCRTcall is trained using a joint loss approach and is enhanced with gated depthwise separable convolution and relative position embeddings. Our experiments demonstrate that GCRTcall achieves state-of-the-art performance in nanopore RNA sequencing signal basecalling, outperforming existing methods in terms of accuracy and robustness. Theses results highlight the effectiveness of integrating advanced transformer architectures with convolutional enhancements for improving RNA sequencing accuracy.

Overall, GCRTcall represents a step forward in nanopore RNA sequencing, offering a robust and precise solution that can facilitate a deeper understanding of transcriptomics and other related fields.

## Acknowledgements

This work is partly supported by grants from the Ministry of Science and Technology of China (2019YFA0707001 and 2021YFF0700201) and the Strategic Priority Research Program of the Chinese Academy of Sciences (XDB37020102).

## Conflict of Interests

Daqian Wang and Jizhong Lou are co-founders and shareholders of Beijing Polyseq Biotech Co. Ltd. Beijing Polyseq Biotech Co. Ltd. and Institute of Biophysics, Chinese Academy of Sciences have filed a patent using materials described in this article.

## Data and code availability

The data used in training, test, and validating GCRTcall are adopted from the published data by Neumann et al. The code, model weights of the model described in this article can be accessed from Github (https://github.com/liqingwen98/GCRTcall).

## Notes

### Competing Interest Statement

The authors have declared no competing interest.

